# Tunnelling nanotubules are druggable mediators of cancer-niche crosstalk

**DOI:** 10.1101/2023.06.20.545732

**Authors:** Sean Hockney, Jess Parker, Babis Tzivelekis, Helen Blair, Kenny Dalgarno, Deepali Pal

## Abstract

Treatment resistance, conferred onto cancer cells largely by the oncogenic niche, remains a clinically unmet need in leukaemia. Tractable and clinically translatable models that mimic cancer-niche crosstalk remain limited, consequently means of clinically drugging microenvironment-driven cancer treatment resistance remain underexplored. Here we develop a prototype bone marrow (BM) like extracellular matrix (ECM), Vitronectin-Alginate-Laminin (VAL), which comprises animal-free components, displays viscoelastic properties like the human BM, and engrafts a range of patient-derived-xenograft acute lymphoblastic leukaemia (PDX-ALL) samples. We discover that following treatment with oxidative stress-inducing apoptotic therapies, such as dexamethasone, ABT-199 and dexamethasone-ABT-199 combination, PDX-ALL cells reach out to MSC via the formation of tunnelling nanotubes (TNT). Nevertheless, we reveal that ALL-VAL-MSC-TNTs are clinically druggable, as they are absent following treatment with CDH2 antagonist ADH-1, a compound well-tolerated in solid cancer Phase I trials. We ultimately expose a triple drug combination of dexamethasone-ABT-199 and ADH-1, with most synergy area (MSA) scores of >30, that shows high efficacy and disrupts functional cancer-niche-TNTs in 4 different high risk PDX-ALL samples. In summary, here we develop prototype cancer-ECM-niche organoids and using leukaemia as a disease paradigm, we provide proof-of-concept insights enabling the beginning of research into drugging functional cancer cell crosstalk with its surrounding cellular and ECM niche.

## Introduction

Cancer treatment resistance remains a clinically unmet need in leukaemia management. Leukaemia and its surrounding BM-niche are in constant crosstalk, especially following therapy, leading both the cancer and its microenvironment to evolve[1]. A key purpose of leukaemia cell interaction with the surrounding niche is to seek protection from treatment, including assaults caused by reactive oxygen species (ROS)-inducing apoptotic agents[2]. Key modes of contact by which leukaemia cells reach out to the surrounding niche cells include direct cell-contact amongst adjacent cells, and for cells in close proximity, via the formation of cytoplasmic projections, known as tunnelling nanotubes (TNT)[3]. Irrespective of the exact mode of contact, cell morphology and polarity of both cancer cells, including leukaemia[4], and niche cells impact these interactions, consequently regulating cell behaviour and function[5]. Nevertheless, means of clinically targeting the malignant niche, including the dynamism of cancer-microenvironment interactions remain unexplored. With this persists the need for stratified oncology driven biomimetic models that enable tracking of real time therapy-induced leukaemia-niche networks.

Next generation organoid models have come a long way in recent times and their importance as human relevant systems with potential to replace animal testing has become increasingly prominent [6]. The importance of such models has gained significant attention since the Food and Drug Administration’s (FDA’s) recent announcement that it will no longer be a requirement for all drugs to be tested on animals prior to in-human trials. Indeed, the FDA itself has now committed to finding new human relevant and non-animal technologies to replace the use of laboratory animals in the development of new drugs and products.

Clinically relevant and human-based animal replacement models are especially warranted in haematological malignancies, where the role of microenvironment in driving treatment resistance is of major significance[7]. Furthermore, transferrable, defined and reproducible co-culture [8], spheroid and organoid [9, 10] models to efficiently capture leukaemia-niche communications are still emerging. Despite significant advances in human cell-based technologies and state-of-the-art organoid platforms [11], a key limitation remains lack of appropriate ECM that are defined in chemical composition, devoid of animal-derived components such as Matrigel, and display physiologically relevant mechanical properties[12]. A key pre-requisite of animal replacement models that would determine their endorsement by the academic and pharmaceutical sector is the tractability of these models. This is to ensure that these models can be meaningfully utilised to study complex biology such as cell-cell crosstalk, cell-cell signalling and other intricate biology, including those involving the ECM.

Here we develop a stratified oncology-driven leukaemia model, and co-culture a range of high risk PDX-ALL samples with BM-ECM-MSC organoids. We apply our model to reveal insights into mechanisms of cancer-niche crosstalk between therapy-primed versus therapy-naïve leukaemia cells and ultimately expose means to directly target such cancer-microenvironment crosstalk using clinically actionable therapies.

## Results

### A prototype animal free, BM-mimicking hydrogel shows high durability to sheer stress, whilst preserving mechanical properties analogous to human BM

The human BM is comprised of both cellular- and extra-cellular matrix (ECM)-components, which are essential for the proper functioning of haematopoietic cells, MSC, osteoblasts and other key cellular constituents of the haematopoietic niche [1, 13–15]. The role of key ECM components, such as laminin and vitronectin in the BM haematopoietic niche is well established [16–22]. To mimic some of the key ECM components *in vitro*, we develop a prototype animal component free, alginate-based hydrogel VAL (Vitronectin-Alginate-Laminin) (Figure 1, Figure S1). We perform ionic cross-linking via calcium chloride, and further enhance the alginate base with rh-laminin-521 and XF-vitronectin. We show effective initiation of cross-linking in VAL (Figure 1B), and over 60% retention of the cross-linked structure over a seven-day period (Figure 1C.). Moreover, we confirm that crosslinked VAL undergoes minimal degradation over a four-day period in both PDX-ALL and MSC tissue culture media, displaying minimal degradation in MSC media over a seven-day period, and retention of at least 50% of initial starting weight in PDX-ALL media (Figure 1D). These data corroborate that crosslinked VAL is stable and is furthermore compatible with cell culture media used for PDX-ALL and MSC culture.

**Figure 1:**
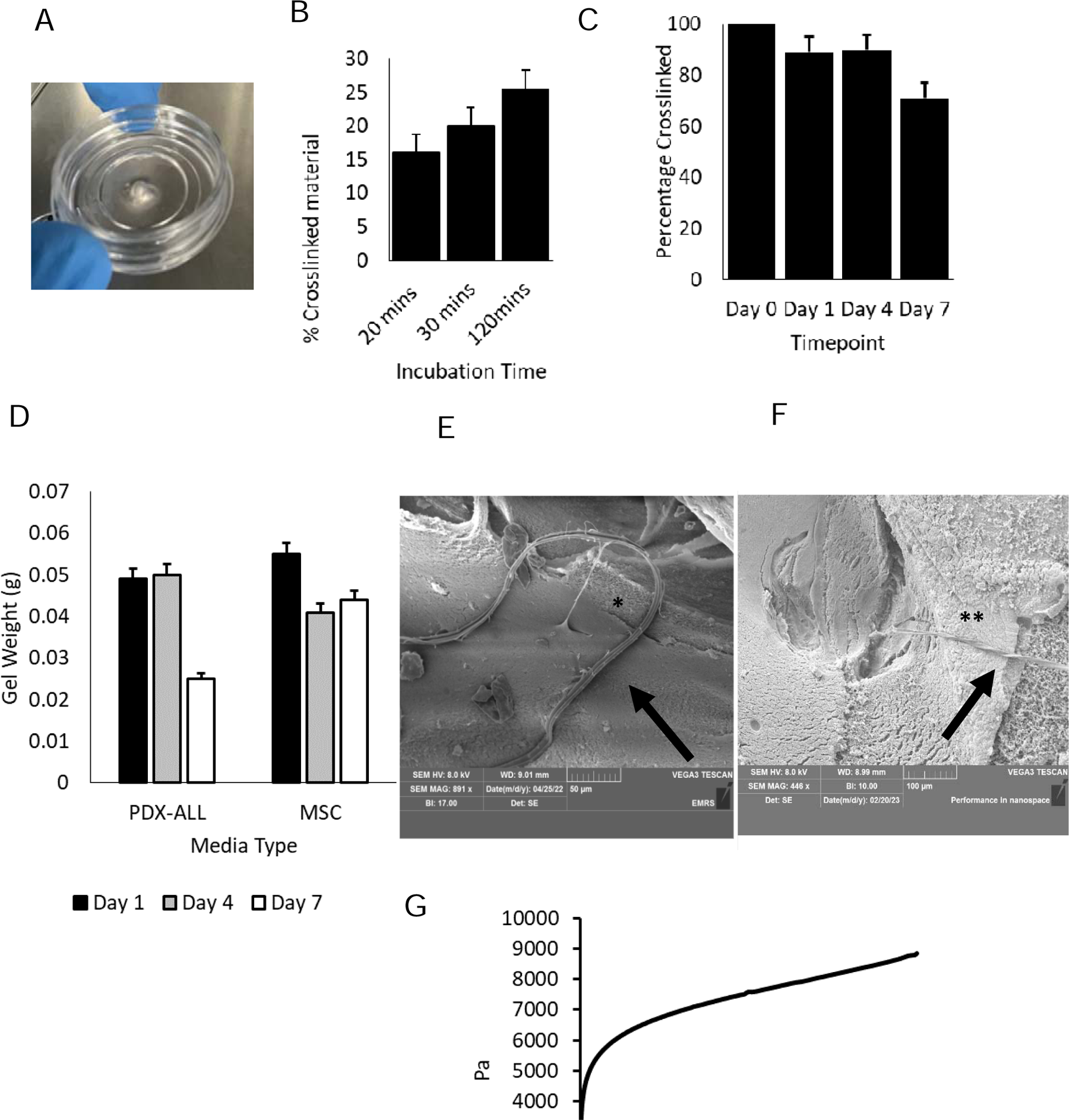
A prototype animal free, BM-mimicking hydrogel shows high durability to sheer stress, whilst preserving mechanical properties analogous to human BM. This figure refers to Supplemental Figure 1. A) Photograph of crosslinked VAL gel. B) % crosslinked VAL at 20, 30 and 120 minutes post crosslinking reaction. Dry, crosslinked material weighed at each timepoint, following removal of excess liquid. C) % corsslinked VAL over a 7-day period. Dry gel material was weighed on each timepoint following removal of any excess liquid. Gels were weighed using a measuring balance D) VAL gel weight measurements following a 7-day exposure to PDX-ALL and iMSC culture media E) Scanning Electron Microscopy (SEM) of laminin (black arrow) bonded via tubule-like structures (*) with alginate in alginate-liminin gel. Image captured at SEM magnification of x882, BI 13, scale bar represents 50µm. F) SEM image showing laminin (black arrow) and vitronectin (**) within VAL biogel. G) Rheological measurements of VAL measuring storage modulus over increasing angular frequency in the range of 0-600 rad/s.

We perform Fourier Transform Infrared Spectroscopy (FTIR) to characterise the chemical bonds within VAL, and find presence of well-established alginate bond types, including C-H, CHO(C=O), COO-, Amide (N-H) and COO-bonds (Figure S1A). We conduct scanning electron microscopy (SEM) experiments, in the first instance, on cross-linked alginate-base hydrogel, which shows a smooth surface (Figure S1B), and then on alginate-base hydrogel infused with laminin, which shows ribbon-like structures (Figure S1C, Figure 1E). We further show laminin matrix incorporation within the cross-linked alginate base, and find the formation of elongated rope-like structures that anchor the laminin within the cross-linked alginate (Figure 1E). Furthermore, we perform additional SEM imaging of the fully formed VAL, and find the presence of textured amorphous particles (Figure 1F), suggesting successful incorporation of vitronectin within VAL.

It is well established that human BMs have viscoelastic properties, which regulate cellular function. Moreover, Young’s modulus, which is a measure of stiffness, and analogous to storage modulus when referring to viscoelastic substances such as the BM, ranges from 250 to nearly 250,000 Pa in human BMs[23]. To define viscoelastic properties of VAL we perform rotational rheological measurements and find that with increasing angular frequency or sheer strain, loss modulus, which measures energy that is lost during the applied stress, remains unchanged (Figure.1G). Moreover, we find that storage modulus, which describes the amount of energy needed to distort the gel, that is, a measure of the stiffness of the gel, increases (Figure.1G). At all times storage modulus remains higher than loss modulus (Figure 1G, Figure S1D), indicating the ability of the gel to resist deformation and retain original physical crosslinks as it returns to its initial state post stress. Furthermore, storage modulus from both the angular frequency- and time-sweep experiments yield values between 4000-9000 Pa (at angular frequency values of 0-600 rad/s), and 3500 Pa achieved at 10 minutes post cross-linking (Figure 1G, Figure S1D). These data substantiate viscoelastic properties of VAL and indicate that the gel retains structure, including preservation of mechanical properties that are comparable with human BM tissues, despite increasing sheer stress.

### VAL shows cytocompatibility with iMSC over a 14-day period

We differentiate human iPSC into MSC, iMSC (Figure S 2A), as per previously published protocols[15], (Figure S 2D) and confirm down regulation of pluripotent genes and upregulation of mesenchymal genes in iMSC (Figure S 2H), and expression of mesenchymal cell surface markers in iMSC (Figure S2E-G). We assess cytocompatibility of VAL, and its ability to support the proliferation of iMSC over 14 days. In the first instance we show SEM images of MSC forming cell aggregates and interacting with VAL (Figure 2A, Figure S 2C). We perform live cell imaging using Hoechst-33342 dye to stain the nucleus and actin to stain the cytoskeleton (Figure 2B), and find that MSC form spheroid-like aggregates within VAL (Figure S 2B). Following a 14-day culture within VAL, we trypsinise VAL-MSC and subsequently extract viable cells. Trypan blue exclusion cell counts at start of cultures (day 0) and then day 1 and day 14 show that VAL sustain starting MSC cell numbers and furthermore, we find nearly two-fold increase in iMSC cell counts when cultured within VAL for 14 days (Figure 2C). In addition, we show that viable iMSC extracted from VAL after 14-day culture retain the expression of key iMSC genes, namely CDH2, NT5E, NES, THY1 and CD44 (Figure S2I) and protein markers, namely ENG and THY1 (Figure 2D). Together these data corroborate the ability of VAL to sustain viable and ENG^+^THY1^+^ iMSC over a 14-day period.

**Figure 2.**
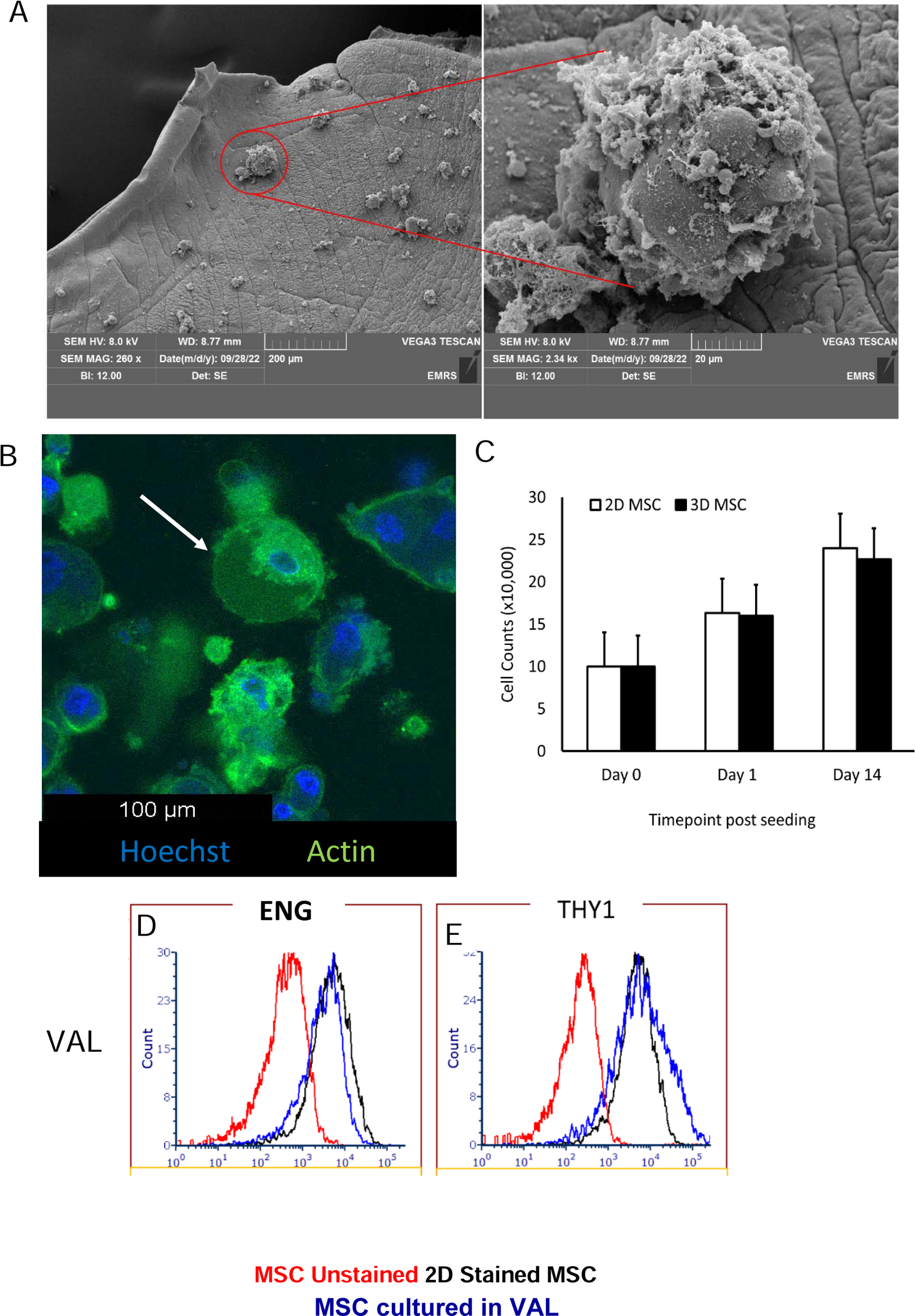
VAL shows cytocompatibility with MSC over a 14-day period. This figure refers to supplemental figure 2. A) Scanning Electron Microscopy of mesenchymal stem cells within the bio-gel where cells group into small 3D clusters. Images taken at 260x magnification and zoomed inlet shows 2.34kx magnification. B) Hoechst actin staining of 3D MSC spheroids within VAL gel shows aggregation of MSC into self-assembling spheroidal structures. C) Cell counts of MSC in 2D and 3D cultures in VAL over a 14-day period. D-E) Flow cytometry data showing fluorescence intensity of live MSC extracted from VAL following 14-day culture, and subsequently stained with cell surface markers D. ENG and E. THY1.

### PDX-ALL primed by VAL upregulate cell-adhesion molecule N-Cadherin (CDH2)

Next we assessed if VAL is able to engineer an oncogenic microenvironment, and consequently engraft and sustain the growth of PDX-ALL over long time periods, that is, in cultures extending beyond 14 days (Figure 3, Figure S3-4). In the first instance, we explore if PDX-ALL cells would engraft, and interact with vitronectin and laminin components within VAL. We test the growth of a total of 5 PDX-ALL samples displaying a range of genetic mutations, namely, 2 high risk BCR::ABL ALL samples, 1 very high risk E2A::HLF ALL sample and the corresponding matched relapse sample containing a homozygous deletion of the glucocorticoid receptor gene NR3C1, and 2 KMT2A-rearranged samples (Figure 3 A-E. Figure S3 A-H). We seed ALL cell suspension from the PDX-ALL samples in cultures surrounding acellular VAL, and in parallel experiments, we seed PDX-ALL within VAL at the time of crosslinking. We find that within 24 hours after seeding, when seeded around VAL, 50% of PDX-ALL migrate to and embed within VAL; and when integrated within VAL at time of crosslinking, then 90% of PDX-ALL remain within VAL (Figure S 4A). In addition, we observe that even after 7 days post seeding, all of PDX-ALL within VAL remain embedded within VAL, and 50% of PDX-ALL seeded around VAL, migrate to and embed within the gel (Figure S4B). Furthermore, we find that seeding PDX-ALL around VAL, enables their migration and engraftment within VAL, which consequently support superior ALL proliferation compared to when the PDX-ALL are encased within VAL at time of culture set up (Figure S4C). These data reveal the chemotactic properties of VAL with respect to leukaemia cell migration, engraftment and retention. Furthermore, we find that ALL cells form aggregates within VAL and these cell aggregates bind to laminin and interact with vitronectin (Figure 3 A-B, Figure S4 A-B). Subsequently we show that PDX-ALL cells proliferate to a greater extent, with reduced time taken to undergo one doubling, in 3D VAL organoid cultures compared to VAL-free suspension cultures (Figure 3 C-E, Figure S4 C-H). In a previous study we had discovered that BM niche primed patient derived leukaemia cells upregulated CDH2, and that CDH2-mediated leukaemia-niche interactions was druggable via ADH-1 [15, 24]. Subsequently, another study found leukaemia cells to acquire mesenchymal gene signatures following interaction with the niche[25]. We investigate the effect of ECM-priming on PDX-ALL and evaluated expression of mesenchymal and anti-apoptotic genes by PDX-ALL following interaction with VAL. We assay the differential gene expression of CDH2, BCL2, MCL1, NR3C1 and NT5E (Figure 3F and Figure S 3I) and find no change in expression of anti-apoptotic genes, including similar expression levels of BCL2 between VAL-primed and VAL-naïve cultures. However, we reveal that PDX-ALL primed by VAL over a 7-day period upregulate the cell adhesion molecule CDH2 (Figure 3F) and the NR3C1 gene which encodes the glucocorticoid receptor (Figure S 3I).

**Figure 3:**
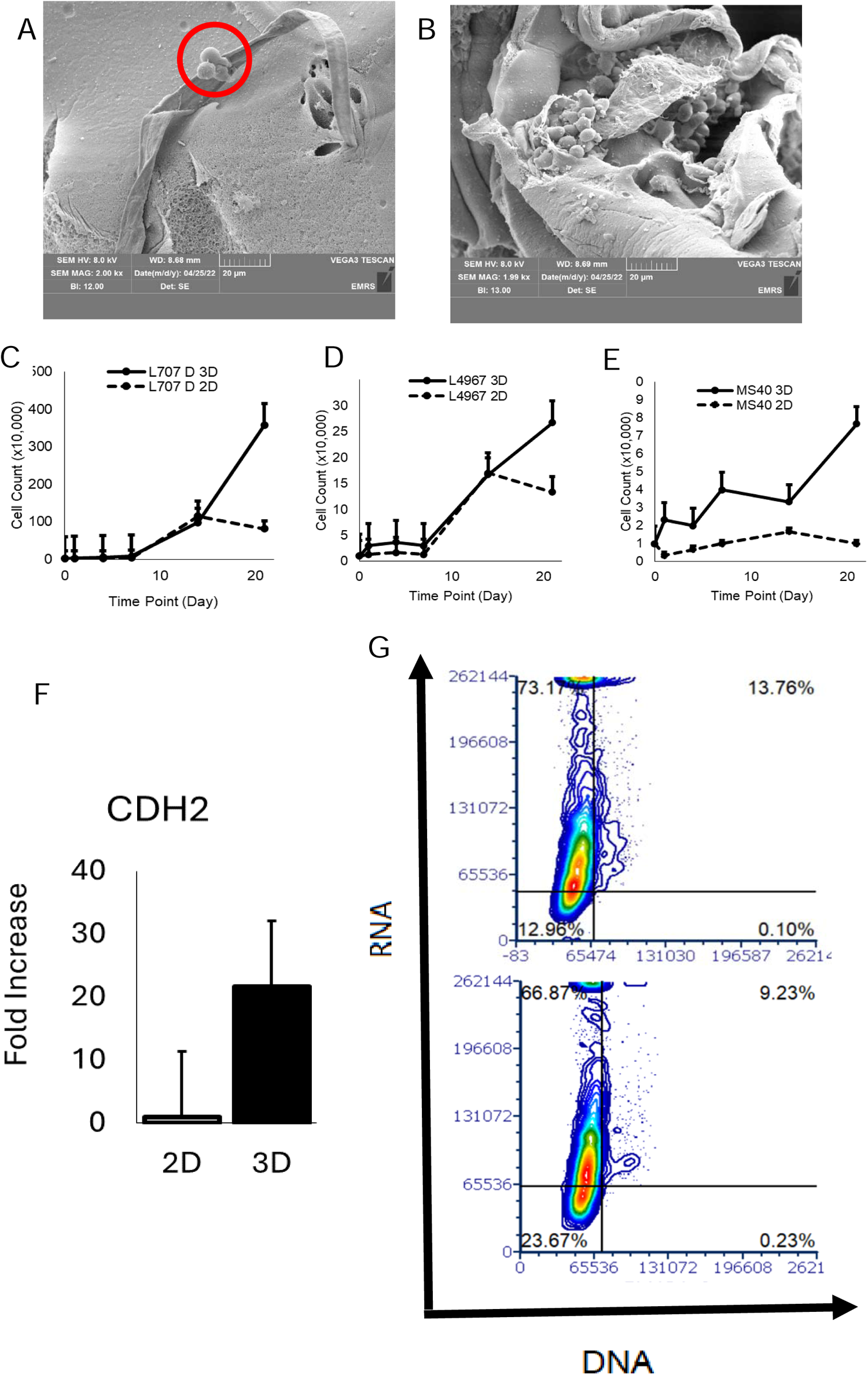
PDX-ALL primed by VAL upregulate cell-adhesion molecule N-Cadherin (CDH2). This figure refers to supplemental figures 3 and 4. A-C) Scanning Electron Microscopy of PDX-ALL incorporated within VAL biogel shows interaction (red circle) between leukaemia, laminin and vitronectin. Images taken at x2.54k (A), x2.00k (B) SEM magnification with BI of 12.00 (A and B). Scale bar= 20µm. C-E) Cell counts of PDX-ALL in 2D suspension cultures versus 3D PDX-ALL-VAL organoids for samples C. L707 carrying E2A::HLF translocation, D. L4967 carrying BCR::ABL1 translocation and E. KMT2A-rearranged mixed lineage (myeloid-lymphoid) sample MS40. SD from N=3. F) qPCR data showing fold change in mRNA expression of CDH2 in 2D versus 3D PDX-ALL-organoid cultures, normalised to GAPDH. G. Contour plot showing DNA (as stained by Hoechst 33342 dye) and RNA (as stained by Pyronin Y in the presence of Hoechst 33342) levels of live patient derived leukaemia cells primed by VAL, over a 5-day period.

Dormant and quiescent cancer cells play a key role in treatment resistance and relapse [13, 15]. However, isolation of patient derived dormant leukaemia cells generally involve mouse serial transplantation assays, which are lengthy experiments and use a lot of animals [13]. We have recently developed protocols to analyse slow cycling and G0 quiescent PDX-ALL in 2D MSC co-cultures[26]. Here we evaluate if VAL can support engraftment of G0, quiescent patient derived leukaemia cells (Figure 3G, Figure S4D-G). We seed PDX-ALL in suspension culture around VAL and following a 5-day culture, harvest VAL-primed PDX-ALL for subsequent G0/G1 analysis. We find that VAL-primed leukaemia cells that did not embed within the ECM possess less than 50% G0 cells compared to PDX-ALL that had successfully embedded within VAL (Figure S4I). Furthermore, we find that under dexamethasone treatment pressure, there are twice as many G0 cells amongst the VAL-embedded PDX-ALL fraction versus the suspension PDX-ALL surrounding VAL (Figure 3G). We repeat these experiments with PDX-ALL obtained from a matched relapse sample, where upon relapse, the patient acquired dexamethasone resistance owing to a homozygous deletion of the NR3C1 gene encoding glucocorticoid receptor. We find similar G0 fractions between PDX-ALL embedded within VAL and PDX-ALL in suspension around VAL to be similar upon dexamethasone treatment. Given the relapse PDX-ALL are insensitive to dexamethasone, these data provide proof-of-concept evidence suggesting the causal association of leukaemia-ECM interactions in mediating leukaemia quiescence. Together these data, corroborate the importance of ECM components in regulating leukaemia cell engraftment and furthermore, G0 and potentially treatment resistance.

### PDX-ALL reach out to VAL-MSC under dexamethasone treatment pressure via the formation of TNTs, which are absent in presence of CDH2 antagonist ADH-1

Reactive oxygen species (ROS) and consequently oxidative stress induced apoptosis is a core mechanism of action of anti-cancer drugs, including targeted therapy agents. MSC interactions with leukaemia cells, including those mediated by TNTs that promote intercellular organelle trafficking, are crucial in protecting malignant cells from treatment-related oxidative stress[3, 27–29]. Capturing these interactions within a 3D ECM-inclusive context is important to recapitulate tissue-relevant cell morphology and polarity, both of which impact cell-cell interactions. We therefore explore mechanisms of leukaemia-MSC crosstalk within PDX-ALL-VAL-MSC co-culture organoid models, in the presence of ROS inducing anti-cancer treatment (Figure 4, Figure S5-6). We observe that in the absence of treatment, PDX-ALL migrate to and attach to MSC within VAL (Figure 4A-C, Figure S5A-C).We find VAL-MSC support viability of PDX-ALL cells (Figure 4C), and moreover proliferation of five high risk PDX-ALL samples with different genetic mutations, namely, 2 BCR::ABL samples, 1 iAMP21 sample, 1 very high risk E2A::HLF sample and 1 KMT2A-rearranged sample. All samples in VAL-MSC co-cultures proliferated at least once every week (Figure S5I),and could be sustained in culture over a period of 21 days (Figure S5D-H). Next we treat the VAL-MSC-PDX-ALL co-cultures with dexamethasone, a glucocorticoid used in clinical management of ALL, where it causes oxidative stress-induced lymphoid apoptosis. We successfully recapitulate clinically proven drug response data, and reproduce both responder and non-responder data, in dexamethasone sensitive and resistant samples respectively (Figure S5K). Moreover, we find PDX-ALL cells co-cultured with VAL-MSC organoids to be less sensitive to dexamethasone than PDX-ALL co-cultured with MSC-free VAL (Figure 4D). Specifically, PDX-ALL co-cultured with VAL-MSC require twice as high a dose to reach < 50% cell viability than PDX-ALL co-cultured with VAL by itself. Since treatment resistance in cancer cells, including leukaemia, have been attributed to TNT-mediated organelle trafficking, we ask if TNTs are being formed in PDX-ALL-VAL-MSC co-cultures, and furthermore if these would be amenable to therapeutic targeting. We begin by investigating mechanistic of TNT formation between PDX-ALL-MSC-VAL organoids, and furthermore their persistence under ROS-induced apoptosis-causing drugs (Figure S6 A-C). We find that in the absence of treatment there are no TNTs being formed between PDX-ALL, MSC and between each other (Figure 4E). However, we reveal that when treated with dexamethasone, then in addition to direct cell contact between MSC-ALL, TNT communication channels begin to form in PDX-ALL-VAL-MSC co-cultures and these are formed between ALL to MSC, ALL to ALL and MSC to MSC (Figure 4E, Figure S6G,I,K). To uncover the effect of apoptosis-causing targeted therapy agents on MSC-ALL TNTs, we repeat these experiments, and this time treat the PDX-ALL-MSC-VAL organoids of BCL2^+^ samples, with the BCL2 inhibitor ABT-199 (Figure 4E, Figure S6D-F). ABT-199 is an oxidative stress-inducing pro-apoptotic agent which is in clinical trials for treating largely high risk, relapsed and treatment refractory ALL. We reveal that TNTs persist under ABT-199 treatment and under treatment with dexamethasone-ABT-199 combination (Figure 4E, Figure S6D-F). Cell-cell communications mediated by the formation of inter-cellular filopodial bridges have been recently suggested to form TNTs, and emerging evidence further alludes for these to be connected by N-cadherin molecules[30]. Given emerging studies, including ours, have found CDH2 to be upregulated by niche-primed PDX-ALL, and here we further discover ECM-primed PDX-ALL to upregulate CDH2, we now ask if blocking N-cadherin would disrupt TNTs in PDX-ALL-VAL-MSC co-cultures. We expose that TNTs disappear after the PDX-ALL-VAL-MSC co-cultures are treated with ADH-1, including in combination with either dexamethasone, ABT-199, and dexamethasone-ABT-199 combination (Figure 4E, Figure SH,J,L). Together these proof-of-concept data suggest that ALL-MSC-ECM crosstalk, in the presence of ROS-inducing therapies, is mediated by TNTs, and PDX-ALL-MSC-ECM TNTs persist following therapy with dexamethasone and ABT-199, both as single and combinatorial agents. We nevertheless expose that these cancer-stroma TNTs are druggable via the FDA orphan drug, CDH2 antagonist ADH-1.

**Figure 4:**
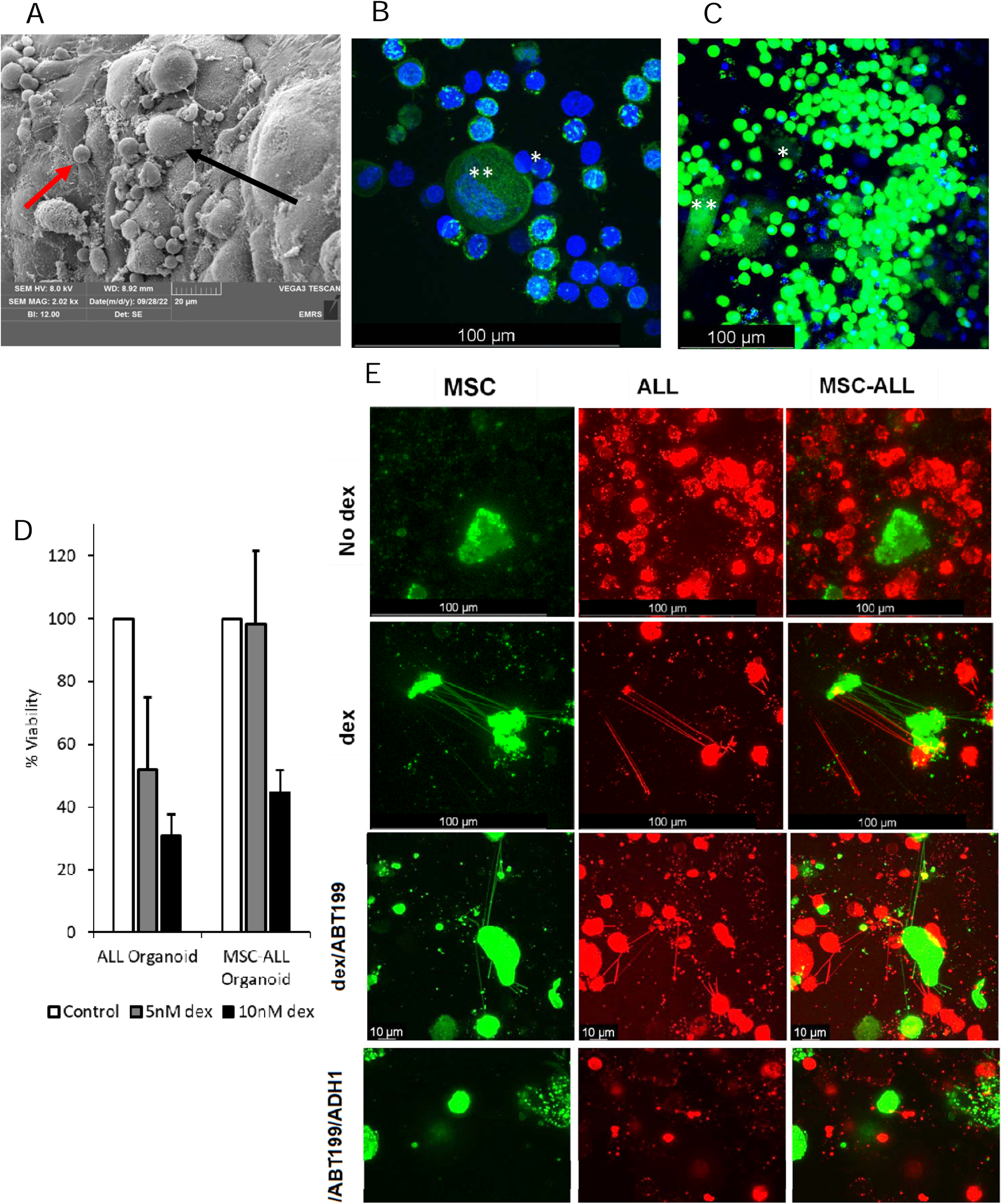
PDX-ALL reach out to VAL-MSC under dexamethasone treatment pressure via the formation of TNTs, which are absent in presence of CDH2 antagonist ADH-1. This figure refers to supplemental figures 5 and 6. A. SEM image showing MSC and leukaemia cells adhering with each other and forming elongated cobweb like inter-cellular tubules. B. Hoechst actin staining of MSC self-assembling spheroids (**) and leukaemia cells (*) within VAL C. Calcein imaging of MSC (**) and patient derived leukaemia (*) cells. D. Trypan blue exclusion cell counts plotted as % viability, of PDX-ALL following dexamethasone treatment in PDX-ALL-VAL-MSC organoids versus PDX-ALL-VAL organoids. E. Live cell confocal imaging of MSC (green) and PDX-ALL sample L707 in VAL-MSC-ALL co-culture. MSC and L707 have been stained with lipophilic tracers DiO and DiL respectively.

### CDH2 antagonist ADH-1 improves efficacy of dexamethasone/ABT-199 in PDX-ALL-VAL-iMSC organoids

ADH-1 has been shown to be well tolerated, with modest efficacy in CDH2 positive solid tumour phase I/II trials, and furthermore received an orphan drug status in 2008 [31]. We have previously shown ADH-1 to be efficacious against PDX-ALL, as a single agent, and in combination with dexamethasone, both in 2D in vitro co-cultures, as well as *in vivo*[8]. Here, we find that PDX-ALL-VAL-MSC co-culture organoids treated with ADH-1, in combination with dexamethasone and ABT-199, yield high efficacy, in three different high risk BCL2 positive PDX-ALL samples (Figure 5A.i., Figure S7A-F). We conduct additional drug combination synergy score analysis on a very high risk E2A::HLF BCL2+ve PDX-ALL sample, where this translocation has proven synergy to dexamethasone-ABT-199 combination. Using SynergyFinder software[32], we apply four different synergy reference models, namely, Bliss, Lowe, Zero interaction potency (ZIP) and Highest single agent (HSA) and reveal synergy scores > 10 across all reference models, for components dexamethasone-ABT-199 and dexamethasone-ADH-1 combinations, and synergy scores > 3 across all reference models, including, Bliss and Lowe, for ABT-199-ADH-1 combination (Fig S7G-I). In addition, we find all three combination components to contain most synergistic area (MSA) scores > 30 (Figure 5A.ii.) at clinically translatable drug concentrations. In summary, we reveal a proof-of-concept triple drug combination, dexamethasone-ABT-199-ADH-1, where individual component drugs are in therapy or have been through Phase I clinical trials, and therefore can realistically be considered for repurposed use in treating high risk leukaemia. Furthermore, we show that in addition to displaying optimal efficacy in PDX-ALL co-cultures with MSC and ECM, this prototype triple drug cocktail, enables the targeting of microenvironment-induced, TNT-driven treatment resistance.

**Figure 5.**
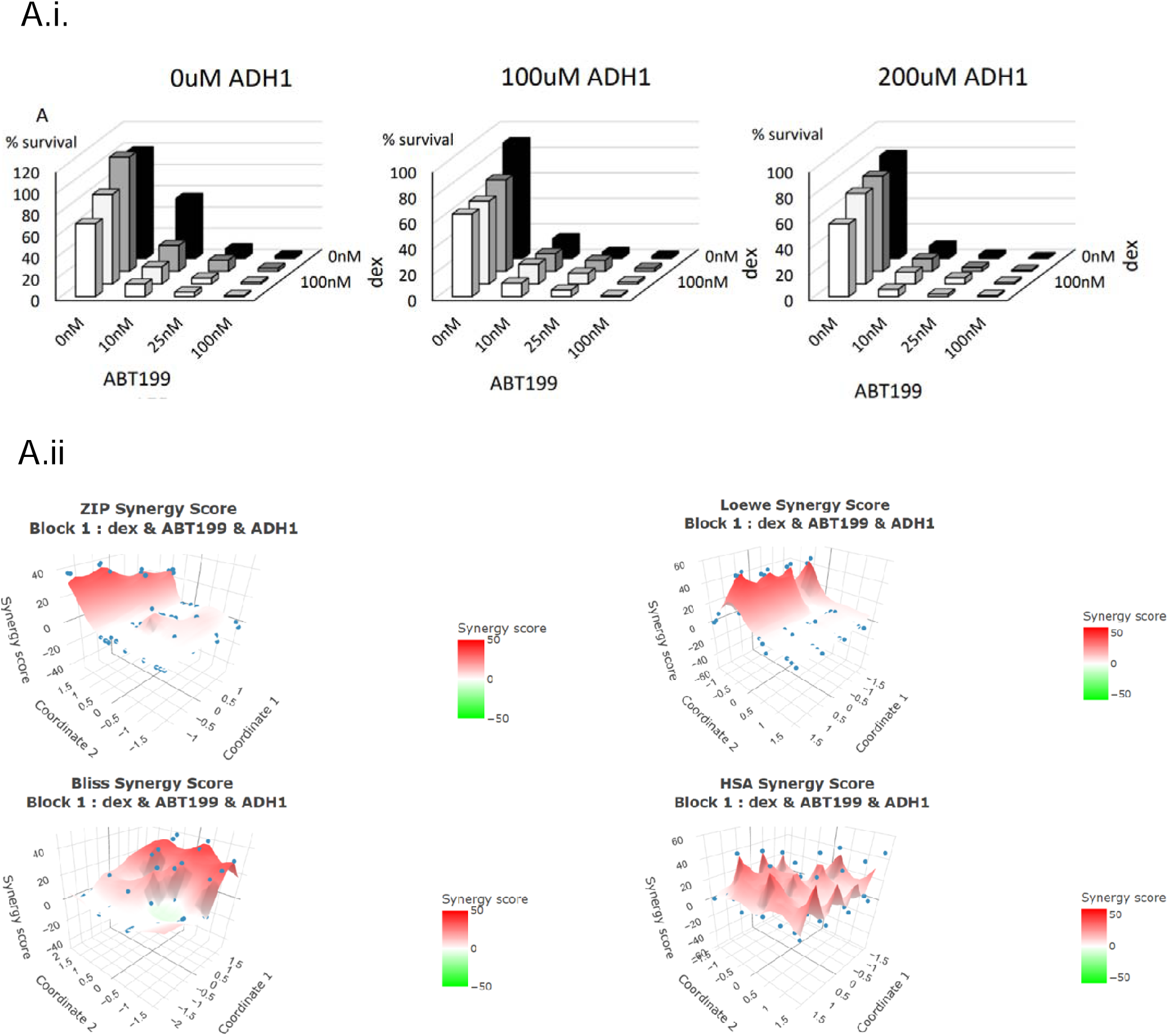
CDH2 antagonist ADH-1 improves efficacy of dexamethasone/ABT-199 in PDX-ALL-VAL-iMSC organoids. This figure refers to supplemental figure 7. CDH2 antagonist ADH-1 improves efficacy of dexamethasone/ABT-199 in PDX-ALL-VAL-MSC organoids. A.i. % Cell survival of PDX-ALL following treatment with dexamethasone-ABT-199 in PDX-ALL-VAL-MSC organoids, with and without the addition of ADH-1. A.ii. ZIP score analysis reveals MSA ZIP scores of > 10 for all three combination arms of dexamethasone-ABT199-ADH-1, namely, dexamethasone-ABT199, dexamethasone-ADH-1 and ABT199-ADH1.

## Discussion

Since 2022, new therapies do not need to be tested in animals to gain FDA approval. On the contrary, there is a paradigm shift in the need for development and use of human tissue relevant, and clinically translatable non-animal technologies, that are tractable, and consequently enable the replacement of animals in preclinical research. Here we develop a prototype animal component-free, chemically defined BM-like synthetic ECM, VAL. We perform in-depth physical, chemical and biological characterisation of VAL and show that it is successfully infused with laminin and vitronectin, key proteins within the human BM microenvironment. We confirm that the viscoelastic properties of VAL, is comparable with that of human BM tissue. Furthermore, we show that VAL is cytocompatible with MSC and is furthermore able to recreate an oncogenic environment, and consequently support engraftment of patient-derived leukaemia cells both individually and in VAL-MSC organoid co-cultures. We apply PDX-ALL-VAL-MSC co-cultures to explore cancer-ECM-MSC interactions, and find that cancer-niche crosstalk is facilitated via both direct cell-cell contact and via formation of TNTs in the presence of treatments.

The role of the malignant BM microenvironment in supporting the survival, growth and treatment resistance of leukaemia is well documented [1, 13–15, 24, 33]. Particularly it is well known that the leukaemia BM niche is dynamic, and its dynamism is reliant on both evolution of leukaemia growth and treatment-driven altered biology [1, 33, 34]. The altered cancer and its niche both deter healthy haematopoiesis and competent immune function in advanced and treatment refractory disease [1, 35].

Several mechanisms have been proposed to be culpable in niche-driven treatment resistance. Stroma signalling via direct cell-cell contact and formation of intercellular crosstalk channels are some key factors mediating leukaemia growth, dormancy and treatment resistance [15, 24, 36]. However, means of clinically targeting such interactions remain obscure. TNTs have recently gained importance as key mediators of cancer-niche communications, particularly their role in protecting cancer cells from ROS induced apoptosis via organelle trafficking [29, 37]. We reveal that in the presence of BM-ECM-components, following treatment with dexamethasone and ABT-199, leukaemia cells reach out to MSC via the formation of TNT. We discover that treatment-induced TNTs can be disrupted by a pentapeptide blocking N-Cadherin. Cadherin adhesion molecules, including cell adhesion molecule N-Cadherin is indeed known to play a role in TNT formation and development [27, 30], and we reveal that therapy-induced, CDH2 mediated niche-leukaemia interactions are amenable to clinical targeting, via ADH-1 [15, 24], a drug that showed a well-accepted toxicity profile in Phase I solid tumour trials [24, 38]. We further expose a highly synergistic triple drug combination to directly target therapy-induced leukaemia-niche TNTs.

Here we show key advances by developing an animal component-free, tissue-relevant synthetic BM-like ECM. To build confidence in application, we implement this cancer-ECM-niche model to discover treatments that directly target leukaemia-niche crosstalk channels, and consequently expose a high synergy triple drug combination. However, the BM microenvironment is inherently complex and besides the mesenchymal niche, it also has sinusoidal and immune niche components. Furthermore, high throughput drug screening platforms require scalable approaches to yield drug dose response data within quick turnaround times. Nevertheless, we provide a prototype to advance the innovation of complex, non-animal, next generation organoids, and we provide proof-of-concept data to enable the beginning of treatment discoveries that directly target functional cancer crosstalk with its microenvironment.

## Methods

### 1. Developing cellular VAL gels

A biogel made of 4% vitronectin and 10% laminin was used along with an alginate base to produce final formulation of VAL biogel. Appropriate volume of VAL mixture added to cultureware and 1:1 volume crosslinking agent (calcium chloride) added to VAL and mixed using gentle pipetting. Biogel was allowed to crosslink at 37°C, 5% CO2 for 20 mins. ALL PDX cells harvested and resuspended at 1×10^6^ cells/ml in fresh culture media. Cell suspension added around biogel following the crosslinking.

### 2. Developing VAL gel MSC-ALL organoids

Biogel as described above added to cultureware. MSC cells harvested to a density of 0.1×10^6^ cells/model. Cells were centrifuged at 1500rpm for 5 minutes to remove media. Pellet was resuspended in equivalent volume of crosslinking agent. MSC-crosslinking agent suspension pipetted into model, and gently mixed to distribute cells evenly throughout model. VAL-MSC models incubated for 20 mins at 37°C, 5% CO2 to crosslink biogel. ALL cells added to crosslinked VAL-MSC models as described in 1.

### 3. DiO-Dil staining of MSC-ALL organoids, treated and untreated

PMSC and ALL harvested from 2D culture at density of 0.1×106 cells/model and 1×106 cells/model respectively. Cells were resuspended at a density of 1×106 cells/ml in fresh serum-free media. 5µl/ml cell suspension of cell labelling solution (DiO for MSC, Dil for ALL) was added and mixed well by gentle pipetting. Cells and staining solution were incubated at 37°C, 5% CO2 for 20 minutes. Cells were then centrifuged at 1500rpm for 5 mins, supernatant removed, pellet resuspended in fresh serum-free media and centrifuged again at same speed and for same time. This process was repeated twice. Following final centrifugation, PMSC and ALL cell pellets were combined and resuspended in volume of crosslinker equivalent to VAL volume. MSC-ALL organoids were left to crosslink for 20 minutes at 37°C and 5% CO2 before fresh culture media was added. Control models had 1ml media added per model, treated models had 1ml media added with 10nM dexamethasone. Models were incubated at 37°C, 5% CO2 for 90 minutes before imaging.

### 4. CellMask Actin Tracking Solution and Hoechst staining of MSC-ALL organoids

PMSC and ALL harvested from 2D culture at density of 0.1×106cells/model and 1×106 cells/model respectively. VAL gel MSC-ALL organoids made as described in 2. 1x CellMask Actin staining solution made from 1000x staining solution using live-cell compatible media. Sufficient 1x CellMask actin solution added to model, enough to cover model. Models were incubated for 60 minutes at 37°C, 5% CO2. CellMask actin solution removed from model and 1x Hoechst solution added, enough volume to cover model. Models incubated for 30 mins before 4 rinsing steps with fresh media. Models imaged.

### 5. Scanning Electron Microscopy of VAL gel models

Samples were fixed in 2% Glutaraldehyde in Sorenson’s Phosphate Buffer overnight before rinsing and dehydration through ethanol series. Samples were dried using a Baltec Critical Point Dryer before mounting onto aluminium stubs with Achesons Silver ElectroDag. Samples were then coated in 15nm of gold using Polaron SEM Coating Unit. Images were captured using a TESCAN VEGA LMU Scanning Electron Microscope housed within the EM Research Services, Newcastle University. Digital images were collected with TESCAN supplied software. All SEM work was completed with assistance from EM Research Services, Newcastle University.

### 6. Rheological assessment of hydrogel formulations

Models were freshly crosslinked directly before rheology analysis as previously described. All experiments were carried out 37°C. Time sweep experiments were carried out over 10 minute and 100 minute time periods. For 10 minute experiments data points were taken every 10 seconds and 100 minute experiments every 20 seconds. Frequency sweep experiments were carried out over a frequency range of 0.1-1000Hz with 500 data points collected within the range. All rheological assessments were completed using an Anton-Paar Rheometer 302e.

### 7. Fourier Transform Infrared Spectroscopy of VAL hydrogel

Samples were crosslinked as described previously immediately before experimentation. Samples were analysed on a Perkin Elmer UATR Two Spectrum Two between wavenumber range of 400-4500cm^-1^, resolution of 4cm^-1^, scan speed= 0.2cm/s. All measurements were obtained with a force gauge of 80.

### 8. Flow cytometry analysis of MSC marker protein expression

Cells were harvested and 1 million cells were taken, washed with PBS and blocked with 10% FBS/PBS. Cells were resuspended in PBS and stained with antibody at 1:40 AB:PBS volume. Samples were incubated at room temperature for 1 hour. Samples were then resuspended in fresh PBS before analysis on FACs CANTO II flow cytometer.

### 9. Hoechst33342/Pyronin Y Cell cycle analysis of PDX-ALL in VAL

Samples were harvested and stained with Hoechst33342 at final working concentration of 10µg/ml. Cells were incubated at 37°C for 45 minutes. At this point 5µl 100µg/ml Pyronin Y was added to each sample before analysis on FACs CANTO II flow cytometer.

### 10. Gene expression analysis via qRT-PCR

RNA was extracted from samples including a DNase I treatment using an RNEasy Micro Kit (Qiagen). cDNA was generated using RevertAIDTMH Minus First Strand cDNA Synthesis Kit. 500ng RNA was used per reaction. The qPCR master mix was composed of 10µM primers, 1x SYBR Green Master Mix and RNAse Free water. qRT-PCR was performed using StepOne Plus RealTime PCR system for 40 cycles including a denaturation step at 95°C, an annealing step at 60°C and an elongation step at 90°C. Analysis was carried out using ΔΔCt fold change method with GAPDH used as housekeeping gene for normalisation.

### 11. Patient derived acute lymphoblastic leukaemia (ALL) cells

Patient-derived xenografts (PDX) were generated as previously described [15]. Male and female 8-12 weeks old mice, immunocompromised NOD.Cg-PrkdcscidIl2rgtm1Wjl/SzJ (NSG) from an in-house colony were transplanted with 1×10^6^ viable patient or PDX ALL cells by intrafemoral injection. Mice were group housed in specific pathogen-free barrier conditions with a 12hour light/dark cycle. PDX ALL blasts cells were isolated from spleen of engrafted humanely euthanized NSG mice when either, high levels of blast were detected by flow cytometry in the peripheral blood or health signs such as anaemia, weight loss and loss of muscle tone were first observed. All regulated animal procedures were approved by the Animal Welfare and Ethical Review Board of Newcastle University and conducted in accordance with the Animals (Scientific Procedures) Act 1996 under the UK Home Office licence P74687DB5.

