## Supplemental file for "Tunnelling nanotubules are druggable mediators of cancer-niche crosstalk"

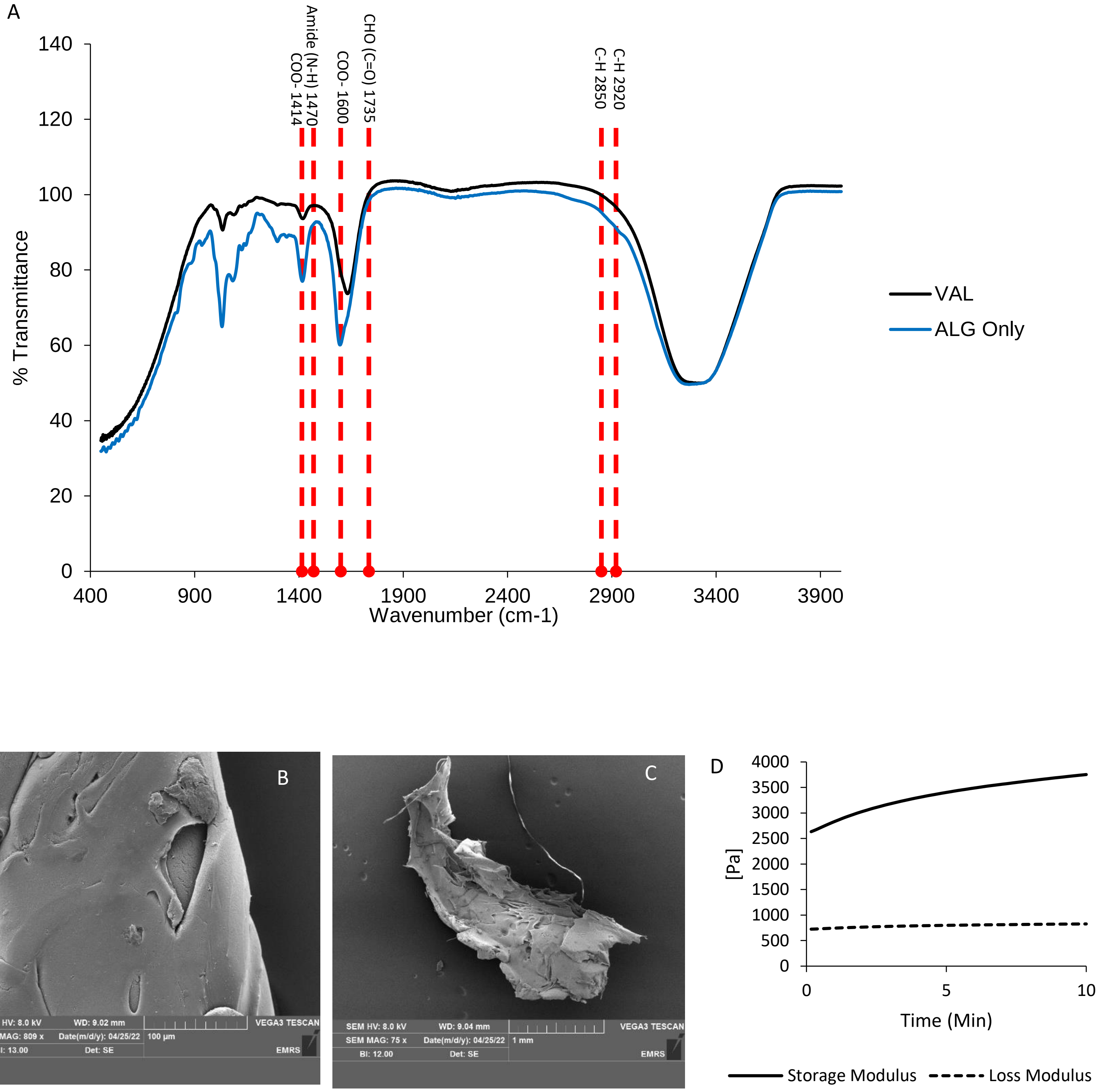
A
% Transmittance
Wavenumber (cm-1)
COO- 1414
Amide (N-H) 1470
COO- 1600
CHO (C=O) 1735
C-H 2850
C-H 2920
VAL
ALG Only
0
20
40
60
80
100
120
140
400
900
1400
1900
2400
2900
3400
3900
B
SEM HV: 8.0 kV
WD: 9.02 mm
SEM MAG: 809 x
Date(m/d/y): 04/25/22
BI: 13.00
Det: SE
100 μm
VEGA3 TESCAN
EMRS
C
SEM HV: 8.0 kV
WD: 9.04 mm
SEM MAG: 75 x
Date(m/d/y): 04/25/22
BI: 12.00
Det: SE
1 mm
VEGA3 TESCAN
EMRS
D
[Pa]
Time (Min)
0
500
1000
1500
2000
2500
3000
3500
4000
0
5
10
Storage Modulus
Loss Modulus

**Supplementary 1 (Relates to Main Figure 1):** Bone marrow ECM components are incorporated into alginate, generating a ECM-laden model. A. Fourier Transform Infrared Spectroscopy of VAL hydrogel and alginate only hydrogel. The presence of well-established alginate bond types are seen and highlighted (1.A) with C-H 2920, C-H 2850, CHO(C=O) 1735, COO- 1600, Amide (N-H) 1470 and COO- 1414 all identified from the spectra for both alginate only and VAL. Subtle differences are seen between the spectra but all key chemical bonds are identifiable for addition of vitronectin and laminin to the alginate. B. SEM image of alginate only hydrogel without BM ECM. The surface is flat and smooth. Scale bar= 100µm, WD= 9.02mm, magnification= 809x. C. SEM of alginate/laminin hydrogel showing ribbons of protein running through the model. Scale bar= 1mm, WD= 9.04mm, magnification= 75x. D. Time sweep rheology of VAL hydrogel over 10 minute period. Viscosity increases over the initial period of crosslinking. At time 0, gel was crosslinked and measurements were taken over the 10 minutes directly following. Storage modulus of VAL increases over time with the loss modulus remaining relatively constant. Storage modulus remains higher than the loss modulus throughout indicating ability of the gel to resist deformation.

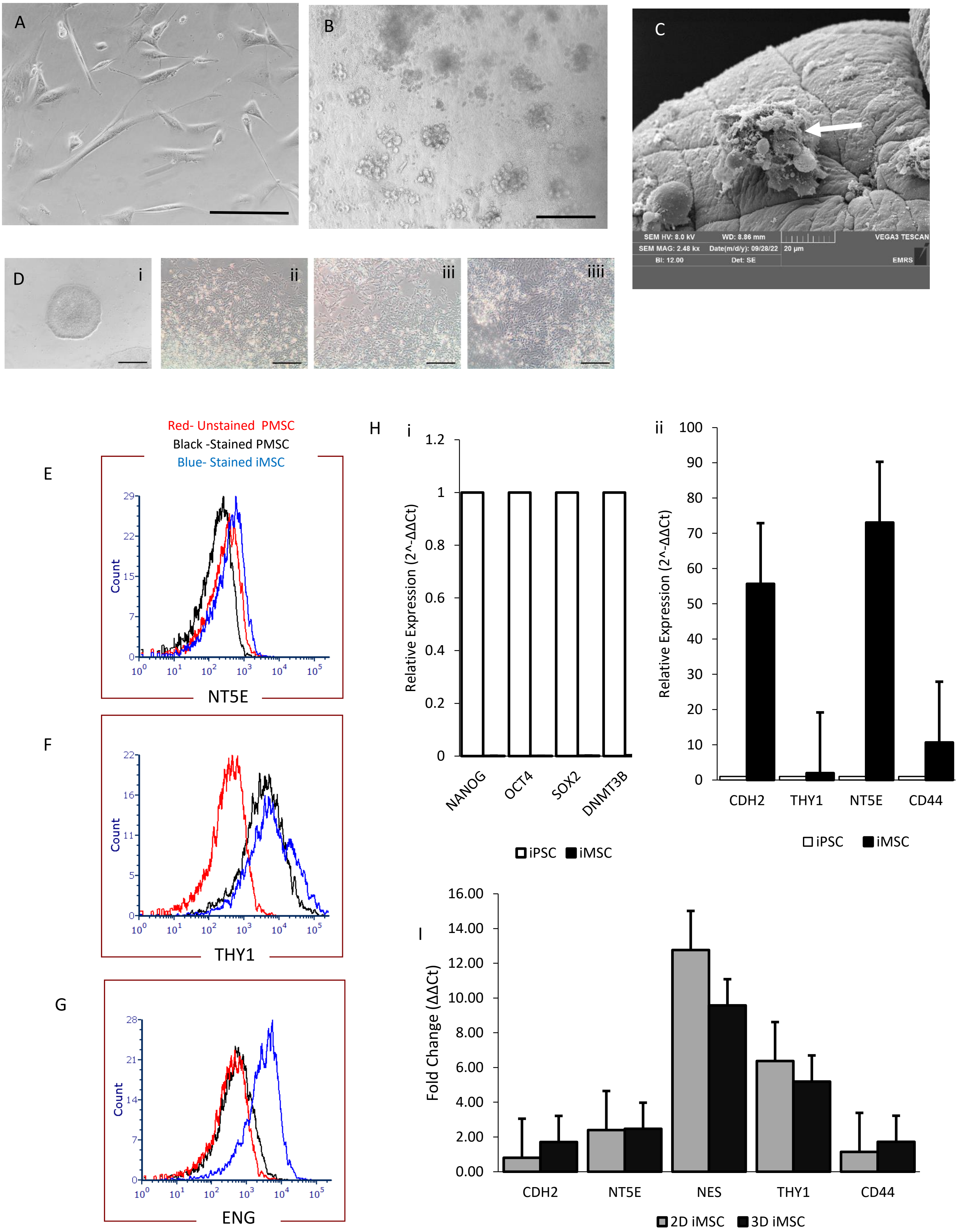
A
B
C
SEM HV: 8.0 kV
WD: 8.86 mm
VEGA3 TESCAN
SEM MAG: 2.48 kx
Date(m/d/y): 09/28/22
20 µm
BI: 12.00
Det: SE
EMRS
D
i
ii
iii
iiii
Red- Unstained PMSC
Black -Stained PMSC
Blue- Stained iMSC
E
Count
NT5E
F
Count
THY1
G
Count
ENG
H
i
Relative Expression (2^-ΔΔCt)
NANOG
OCT4
SOX2
DNMT3B
iPSC iMSC
ii
Relative Expression (2^-ΔΔCt)
CDH2
THY1
NT5E
CD44
iPSC iMSC
I
Fold Change (ΔΔCt)
CDH2
NT5E
NES
THY1
CD44
2D iMSC 3D iMSC

**Supplementary 2 (Relates to Main Figure 2):** Mesenchymal stem cells form spheroid-like structures within VAL hydrogel and maintain MSC marker expression. A. Phase contrast image of 2D MSC, x40 magnification, scale bar= 100µm. B. Phase contrast image of 3D MSC within VAL hydrogel. Spheroid structures form with collections of MSC (white arrow). Magnification=x20, scale bar= 100µm. C) SEM image of MSC spheroid-like structures in VAL hydrogel. Scale bar= 20µm, WD 8.86mm, magnification 2.48kx. D. Microscope images of iPSC-iMSC differentiation via mesoderm induction. Images captured at Day 0 (i), 24 hours (ii), 72 hours (iii) and Day 14 (iiii). Scale bars= 100µm. E-G. Flow cytometry analysis of mesenchymal stem cell markers comparing primary MSC and iPSC-derived MSC. CD73 (E) expression was comparable between PMSC (black peak) and iMSC (Blue peak) with only slight positive expression seen. F. THY1 expression was comparable between PMSC (black peak) and iMSC (blue peak). G. iMSC (blue peak) showed much higher levels of CD105 expression compared to PMSC (black peak) H. qPCR gene expression characterisation of iPSC-derived MSC shows positive expression of MSC-positive markers CDH2, THY1, NT5E and CD44 whilst also showing downregulation of classical iPSC markers NANOG, OCT4, SOX2 and DNMT3B. Relative expression calculated by normalising to GAPDH and fold expression was compared to iPSC expression. N=3. I. qPCR data showing key MSC marker expression pre and post VAL culture after 5 days. CD44 and CDH2 expression was upregulated in 3D MSCs compared to 2D MSC with comparable expression of NT5E between 2D and 3D. NES and THY1 expression was shown to be downregulated in 3D compared to 2D.  N=3.

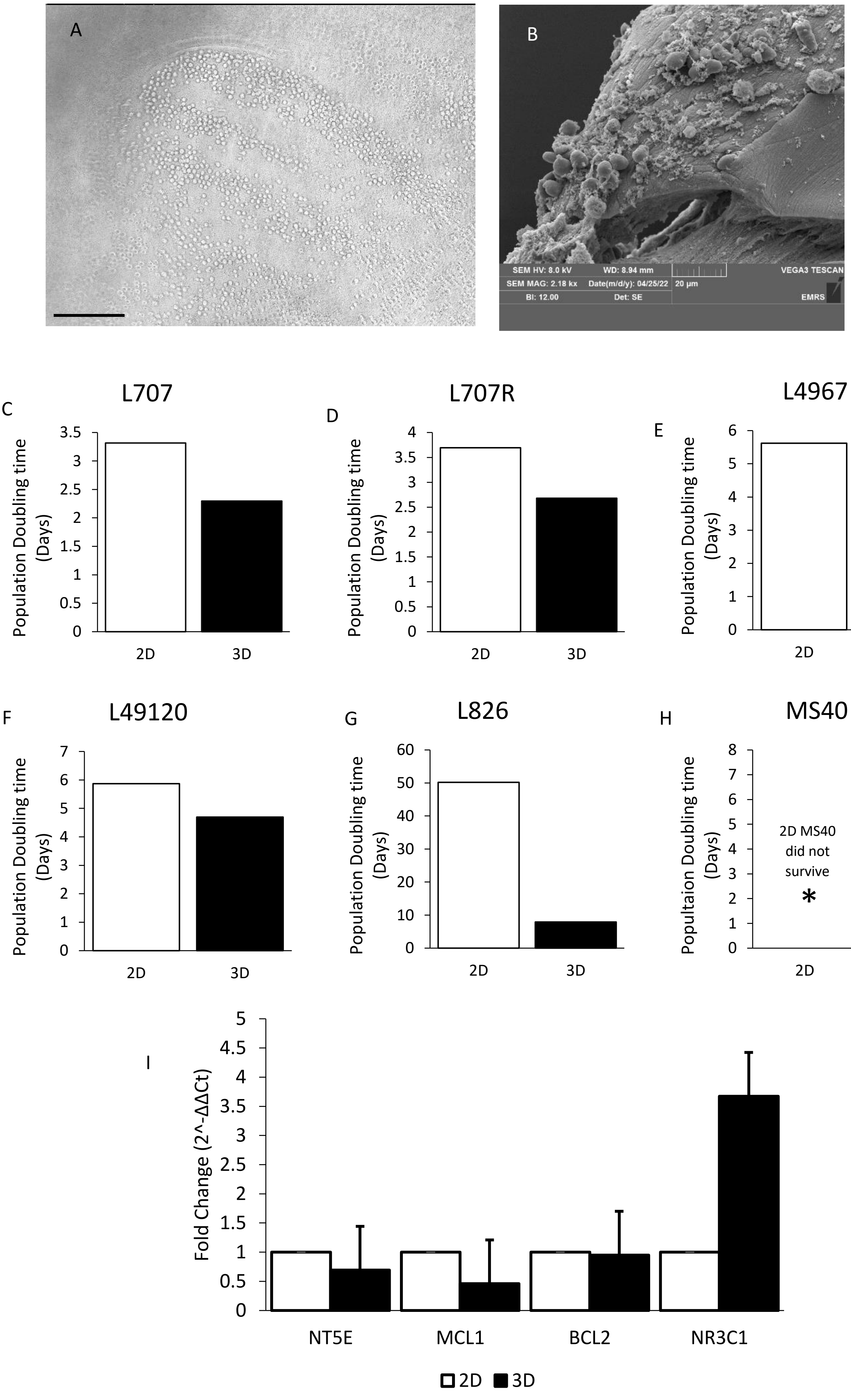

A
B
SEM HV: 8.0 kV
WD: 8.94 mm
VEGA3 TESCAN
SEM MAG: 2.18 kx
Date(m/d/y): 04/25/22
20 µm
BI: 12.00
Det: SE
EMRS
C
L707
D
L707R
E
L4967
F
L49120
G
L826
H
MS40
Population Doubling time (Days)
Popultaion Doubling time (Days)
2D MS40 did not survive
*
2D
3D
I
Fold Change (2^-ΔΔCt)
NT5E
MCL1
BCL2
NR3C1
2D
3D

**Supplementary 3 (Relates to Main Figure 3):** Leukaemia cells embed into VAL hydrogel. A. Phase contrast image of L707 cells in 3D culture with VAL hydrogel. Magnification x10, scale bar= 200µm. B. SEM image of L707 cells embedded into VAL hydrogel and interacting with vitronectin. SEM Magnification= 2.18kx, WD= 8.94mm. Scale bar= 20µm. C-H. Doubling times of PDX samples in 2D monoculture and in 3D VAL culture. In all cases doubling time is reduced in 3D VAL cultures (L707 from 3.3 days in 2D to 2.3 days in 3D; L707R from 3.7 days in 2D to 2.7 days in 3D; L4967 from 5.6 days in 2D to 4.4 days in 3D; L49120 from 5.9 days in 2D to 4.7 days in 3D; L826 from 50.2 days in 2D to 7.9 days in 3D). Note 2D MS40 had no doubling time due to cells not surviving in 2D monoculture (*) with 3D doubling time being 7.1 days. I. qPCR data for ALL marker expression in 2D and 3D L707s. NT5E and MCL1 are downregulated in 3D VAL L707 cells compared to 2D equivalent cultures. BCL2 expression is comparable between 2D and 3D cultured L707 cells. NR3C1 expression is upregulated 3.67x more in 3D VAL cultured L707 cells compared to 2D cultured cells. N=3.

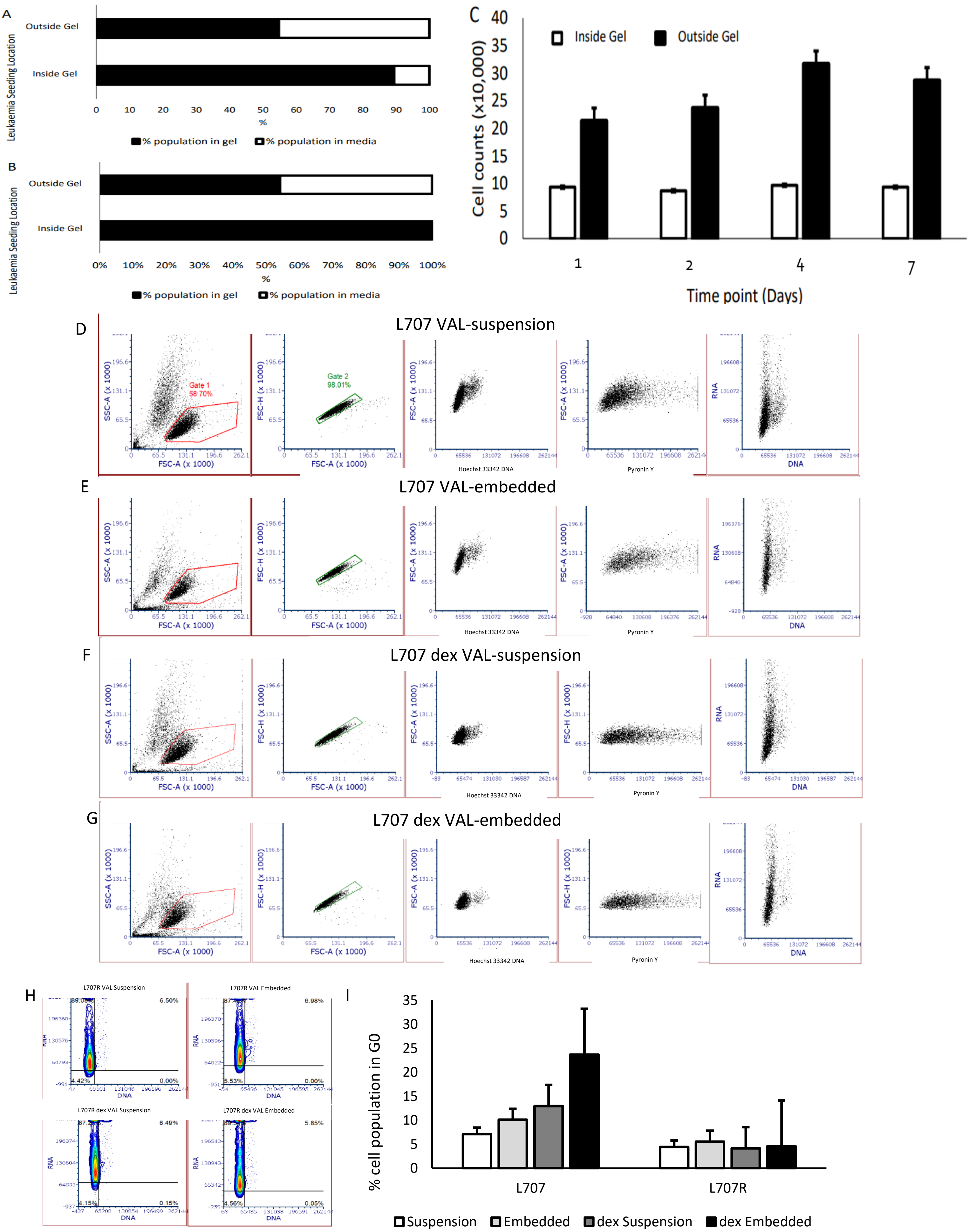

A
Outside Gel
Inside Gel
Leukaemia Seeding Location
0 10 20 30 40 50 60 70 80 90 100
%
% population in gel
% population in media
B
Outside Gel
Inside Gel
Leukaemia Seeding Location
0% 10% 20% 30% 40% 50% 60% 70% 80% 90% 100%
%
% population in gel
% population in media
C
Inside Gel
Outside Gel
Cell counts (x10,000)
Time point (Days)
D
L707 VAL-suspension
Gate 1 58.70%
Gate 2 98.01%
SSC-A (x 1000)
FSC-A (x 1000)
FSC-H (x 1000)
Hoechst 33342 DNA
Pyronin Y
RNA
DNA
E
L707 VAL-embedded
F
L707 dex VAL-suspension
G
L707 dex VAL-embedded
H
L707R VAL Suspension
L707R VAL Embedded
L707R dex VAL Suspension
L707R dex VAL Embedded
I
% cell population in G0
L707
L707R
Suspension
Embedded
dex Suspension
dex Embedded

**Supplementary 4 (Relates to main Figure 3):** A and B. Distribution of PDX following seeding in different locations of the hydrogel model- inside the gel and surrounding the gel. Cell counts were taken at regular timepoints up to 7 days of culture, with cells harvested from each compartment of the model. A. 24hours post seeding we see when cells are seeded outside the gel, approximately 50% of the cells total population have migrated into the model, whereas the majority of cells seeded within in the gel stay within the gel (90%). This trend is seen after 7 days also (B). C. Raw cell counts of populations found within each hydrogel model compartment at Days 1, 2 4 and 7 post seeding. Cells harvested from models seeded outside the gel and allowed to move to and from the model showed increased proliferation over cells that were seeded within the gel. D-G. Dot plots and flow cytometry analysis of Hoechst/Pyronin staining of L707 (matched with the contour plots in Main Figure 3G) with additional analysis for control suspension and embedded cells. H. Contour plots for L707R with Hoechst/ Pyronin staining flow cytometry shows no change in percentage of dormant cells before and after dexamethasone treatment and also no change is seen between VAL suspension versus VAL embedded cell fractions. I. %G0 PDX-ALL between samples taken from a patient at diagnosis (L707) and at relapse (L707R), untreated and following treatment with dexamethasone. At relapse the patient acquired resistance to dexamethasone owing to a homozygous deletion of NR3C1 encoding the glucocorticoid receptor.

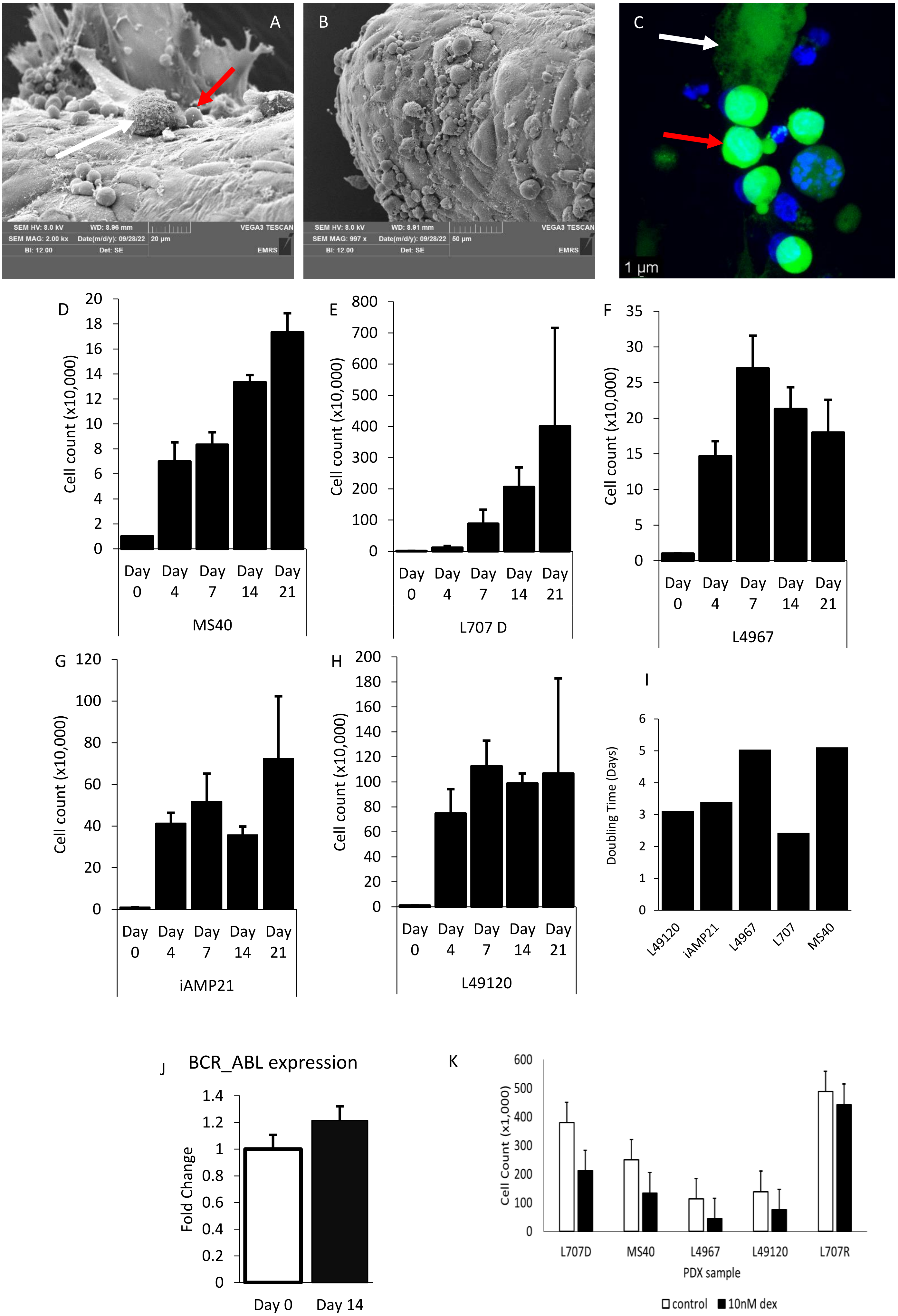
A
SEM HV: 8.0 kV
WD: 8.96 mm
VEGA3 TESCAN
SEM MAG: 2.00 kx
Date(m/d/y): 09/28/22
20 μm
BI: 12.00
Det: SE
EMRS
B
SEM HV: 8.0 kV
WD: 8.91 mm
VEGA3 TESCAN
SEM MAG: 997 x
Date(m/d/y): 09/28/22
50 μm
BI: 12.00
Det: SE
EMRS
C
1 μm
D
Cell count (x10,000)
Day 0
Day 4
Day 7
Day 14
Day 21
MS40
E
Cell count (x10,000)
L707 D
F
Cell count (x10,000)
L4967
G
Cell count (x10,000)
iAMP21
H
Cell count (x10,000)
L49120
I
Doubling Time (Days)
L49120
iAMP21
L4967
L707
MS40
J
BCR_ABL expression
Fold Change
Day 0
Day 14
K
Cell Count (x1,000)
L707D
MS40
L4967
L49120
L707R
PDX sample
control
10nM dex

**Supplementary 5 (Relates to Main Figure 4):** A and B. SEM images of MSC-ALL-VAL co-culture model. Larger MSCs (white arrow) and ALL cells (red arrow) come into close proximity within the 3D model. Scale bars= 20 and 50µm respectively. C. Hoechst calcein image staining of PDX-ALL-MSC co culture at Day 7 of culture, iMSCs (white arrow) and PDX-ALL (red arrow) come into close proximity to each other and demonstrate viability. Magnification= x63, zoom= 2.23x. Scale bar= 1µm. D-H. Long term co-culture proliferation of PDX samples with different cytogenetics. Counts were taken periodically over a 21-day period. I. Population doubling times of PDX samples within the 3D MSC-PDX-VAL co-culture model. All samples had doubling times between 2 and 5 days within the model. J qPCR data from L4967 cells cultured in 3D VAL model shows maintenance of the BCR::ABL translocation following 14 days of culture within the co-culture VAL-ALL-MSC model. N=3. K-L. MSC-PDX-ALL co-culture model can provide clinically translatable drug response data to single agent dexamethasone. L707D, MS40, L4967 and L49120 are known responders to dexamethasone from clinical data and here we show a significant reduction in cell survival in all samples within our in vitro model. L707R is the relapse sample from L707D and developed a known resistance to dexamethasone due to the deletion of the glucocorticoid receptor NR3C1, here we show no response to dexamethasone in this sample within our model. K is % survival and L is raw cell counts. N=3.

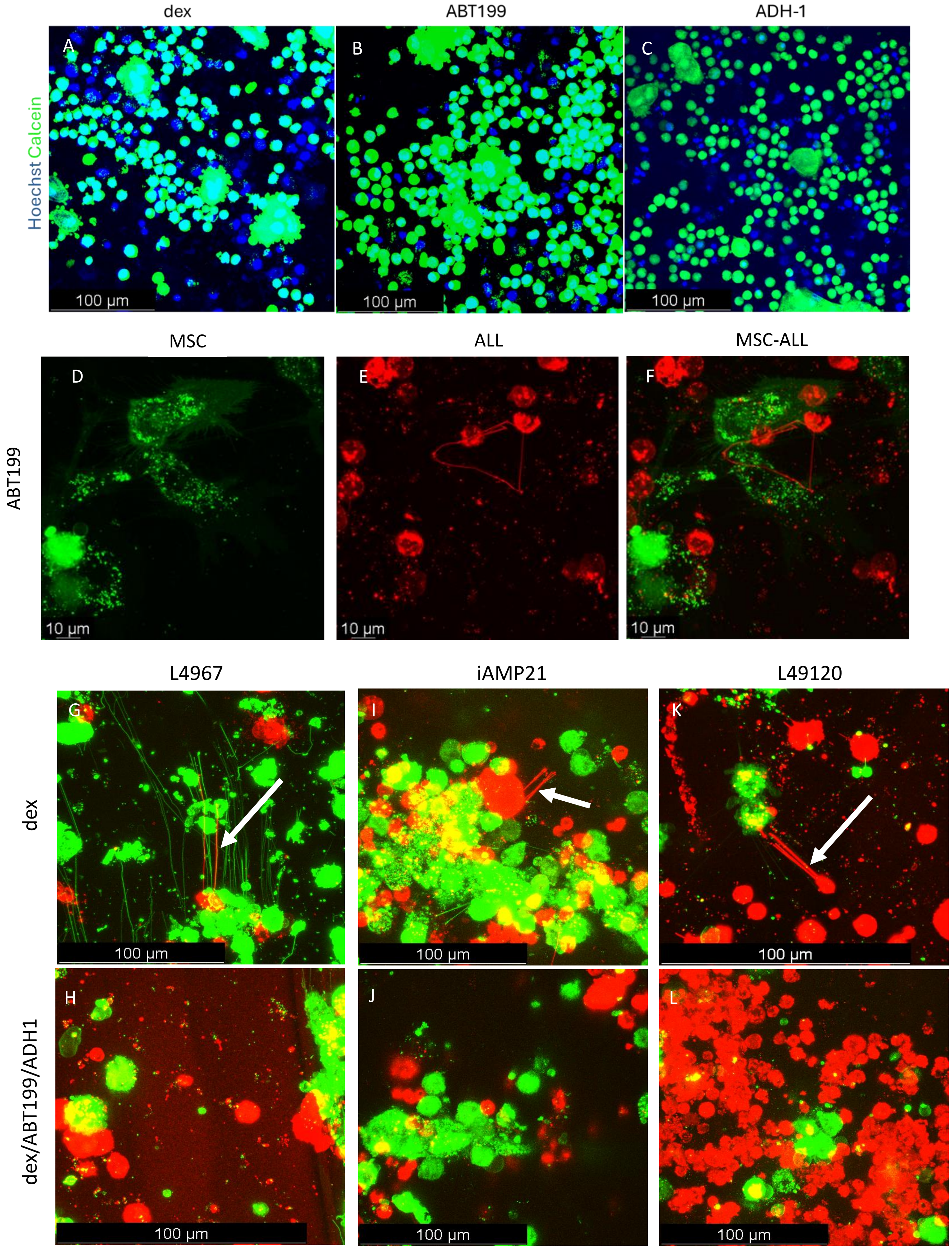
dex
ABT199
ADH-1
Hoechst Calcein
A
B
C
100 µm
100 µm
100 µm
MSC
ALL
MSC-ALL
ABT199
D
E
F
10 µm
10 µm
10 µm
L4967
iAMP21
L49120
dex
G
I
K
100 µm
100 µm
100 µm
dex/ABT199/ADH1
H
J
L
100 µm
100 µm
100 µm

**Supplementary Figure 6 (Relates to Main Figure 4):** A-C. Hoechst/calcein staining of MSC-ALL-VAL co-cultures following 5 days of treatment with 10nM dexamethasone (A), 100µM ADH-1 (B) and 150nM ABT199 (C). Least amount of viable cells is seen following 100µM ADH-1. X40 magnification in oil immersion. Scale bars= 100µm D. DiO/Dil staining of MSC-L707-VAL co-culture models following treatment with 150nM ABT199. TNTs still form from ALL cells (*). G-L. DiO/Dil staining of L4967, iAMP21 and L49120 PDX samples shows that TNTs form from ALL cells under dexamethasone treatment pressure. These TNTs are not present when ADH-1 is present within the treatment combination in the form of dex/ABT199/ADH-1 combination. All TNT imaging experiments were captured in the same way: z stacks of 100µm captured at x 40 magnification in oil immersion with minimum of 20 images captured/ z stack. For each model 5 standardised locations were captured, representative images shown here. All images captured using confocal microscopy (Leica) and analysed via LASX software.

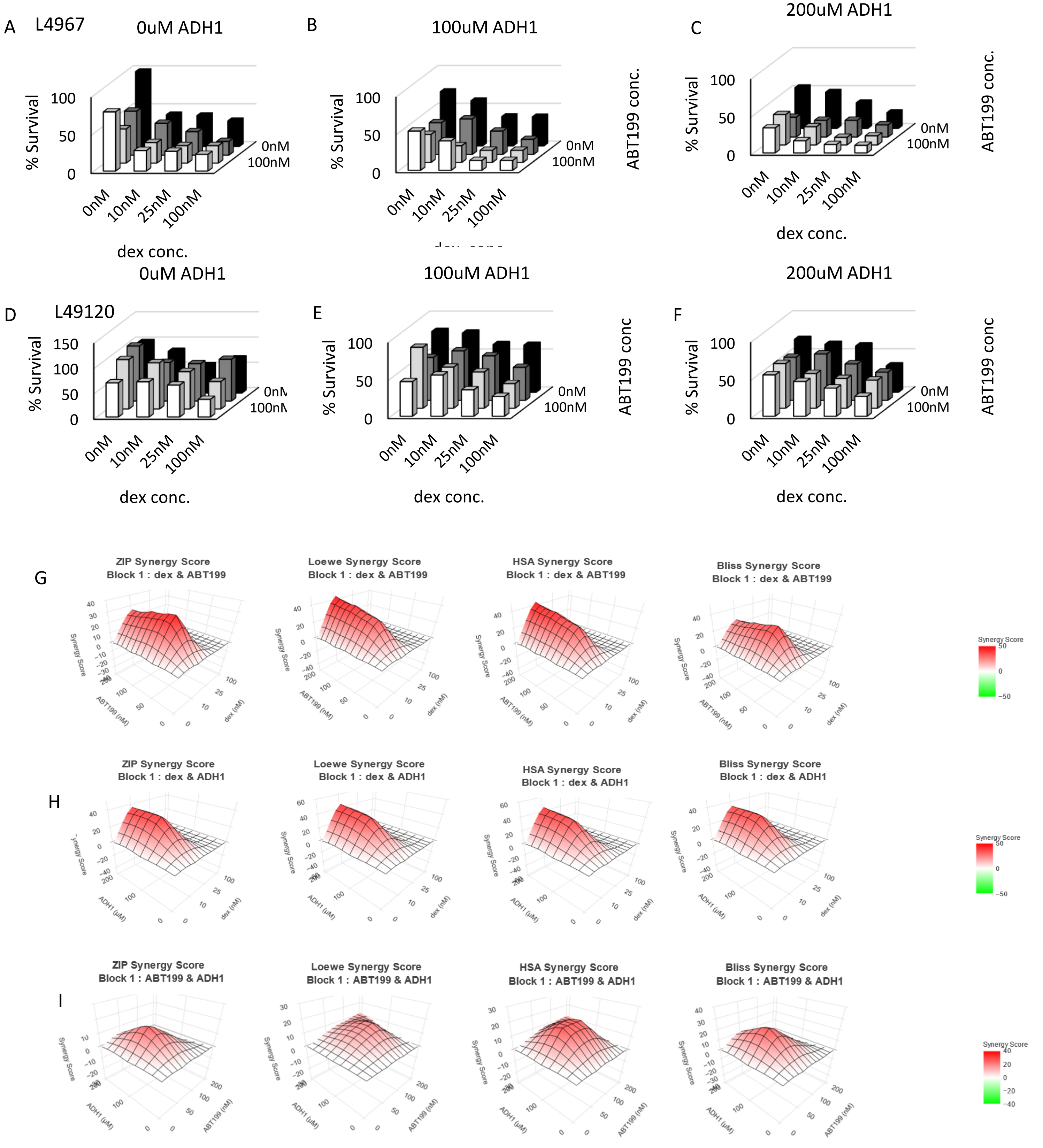
A
L4967
0uM ADH1
% Survival
100
50
0
0nM
10nM
25nM
100nM
dex conc.
0nM
100nM
B
100uM ADH1
ABT199 conc.
C
200uM ADH1
D
L49120
0uM ADH1
150
E
100uM ADH1
ABT199 conc
F
200uM ADH1
G
ZIP Synergy Score
Block 1 : dex & ABT199
Loewe Synergy Score
Block 1 : dex & ABT199
HSA Synergy Score
Block 1 : dex & ABT199
Bliss Synergy Score
Block 1 : dex & ABT199
Synergy Score
ABT199 (nM)
dex (nM)
H
ZIP Synergy Score
Block 1 : dex & ADH1
Loewe Synergy Score
Block 1 : dex & ADH1
HSA Synergy Score
Block 1 : dex & ADH1
Bliss Synergy Score
Block 1 : dex & ADH1
ADH1 (uM)
I
ZIP Synergy Score
Block 1 : ABT199 & ADH1
Loewe Synergy Score
Block 1 : ABT199 & ADH1
HSA Synergy Score
Block 1 : ABT199 & ADH1
Bliss Synergy Score
Block 1 : ABT199 & ADH1

**Supplementary Figure 7 (Relates to Main Figure 5):** dexamethasone/ABT199/ADH-1 combination matrices for L4967 (A-C) and L49120 (D-F). % Cell survival of patient derived ALL samples L4967 and L49120 following dex/ABT199 treatment with and without ADH-1. G-I Synergy score analysis of double components within dexamethasone/ABT199/ADH-1 combination on L707 sample. High Synergy scores are seen using all four Synergy approaches: ZIP, Loewe, HSA and Bliss denoted by red areas. Dex/ABT-199, dex/ADH-1 and ABT-199/ADH-1 combinations all show synergistic relationships.
